## Supplementary information for "Mitochondrial complex I deficiency stratifies idiopathic Parkinson’s disease"

Irene H Flønes *et al*.

* Corresponding author: Charalampos Tzoulis.

**This PDF file includes:**

Figs. S1 to S13

Reference [60]

**Other Supplementary Information for this manuscript include the following (separate files):**

Supplementary Data 1 to 13

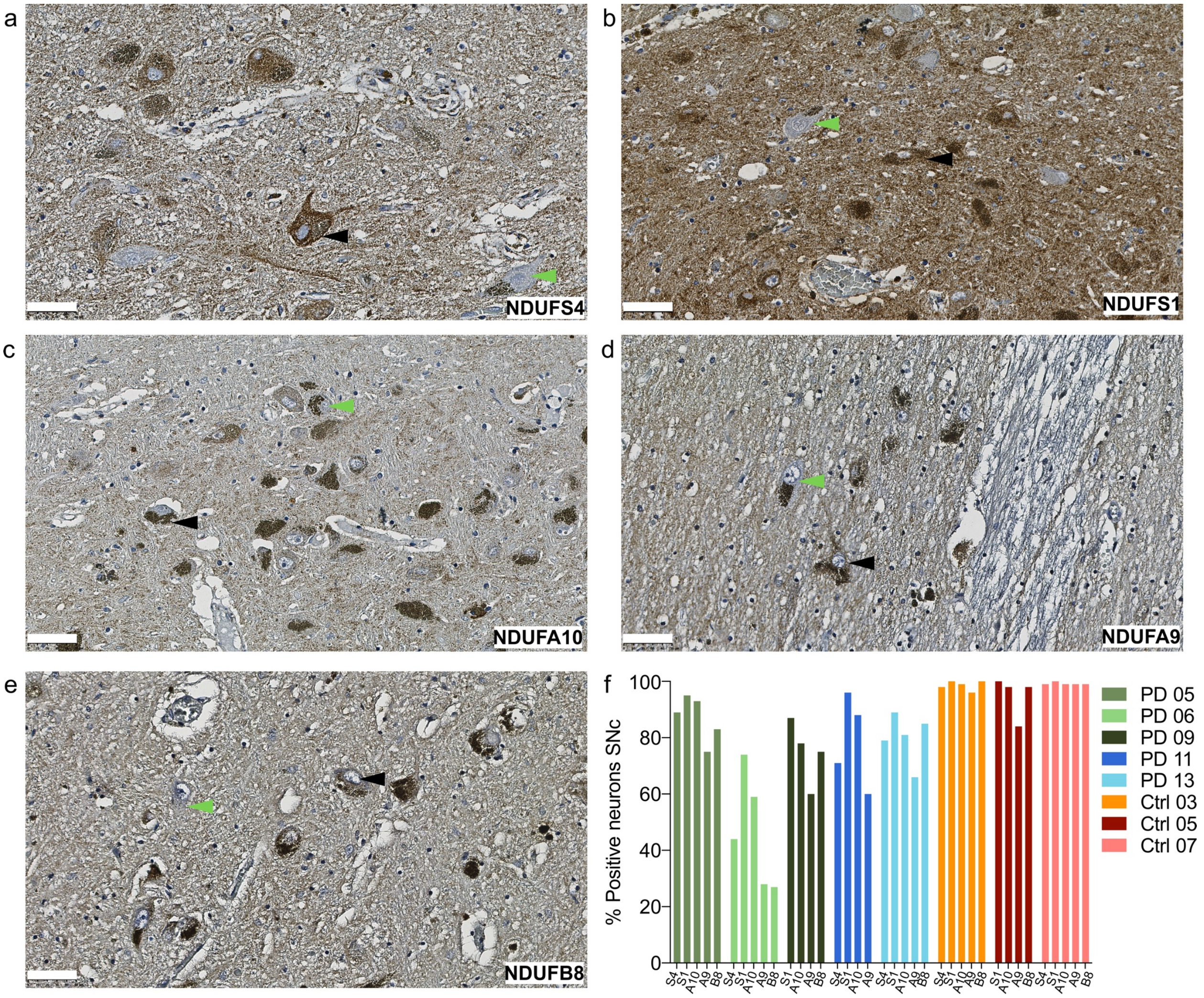


**Fig. S1. Immunostaining of CI subunits in the SNpc**

(a-e) Representative images of immunostaining for CI subunits: NDUFS4, NDUFS1, NDUFA10, NDUFA9 and NDUFB8 in the SNpc of an individual with iPD. Green arrowheads show CI deficient neuromelanin positive neurons. Black arrowheads show CI intact neurons. Scale bar 50 µm. (f) Graph shows the proportion (in %) of CI positive neuromelanin positive neurons (y-axis) in the SNpc of iPD subjects ($n = 5$) and controls ($n = 3$) stained for five different complex I subunits (x-axis). S4: NDUFS4, S1: NDUFS1, A10: NDUFA10, A9: NDUFA9, B8: NDUFB8

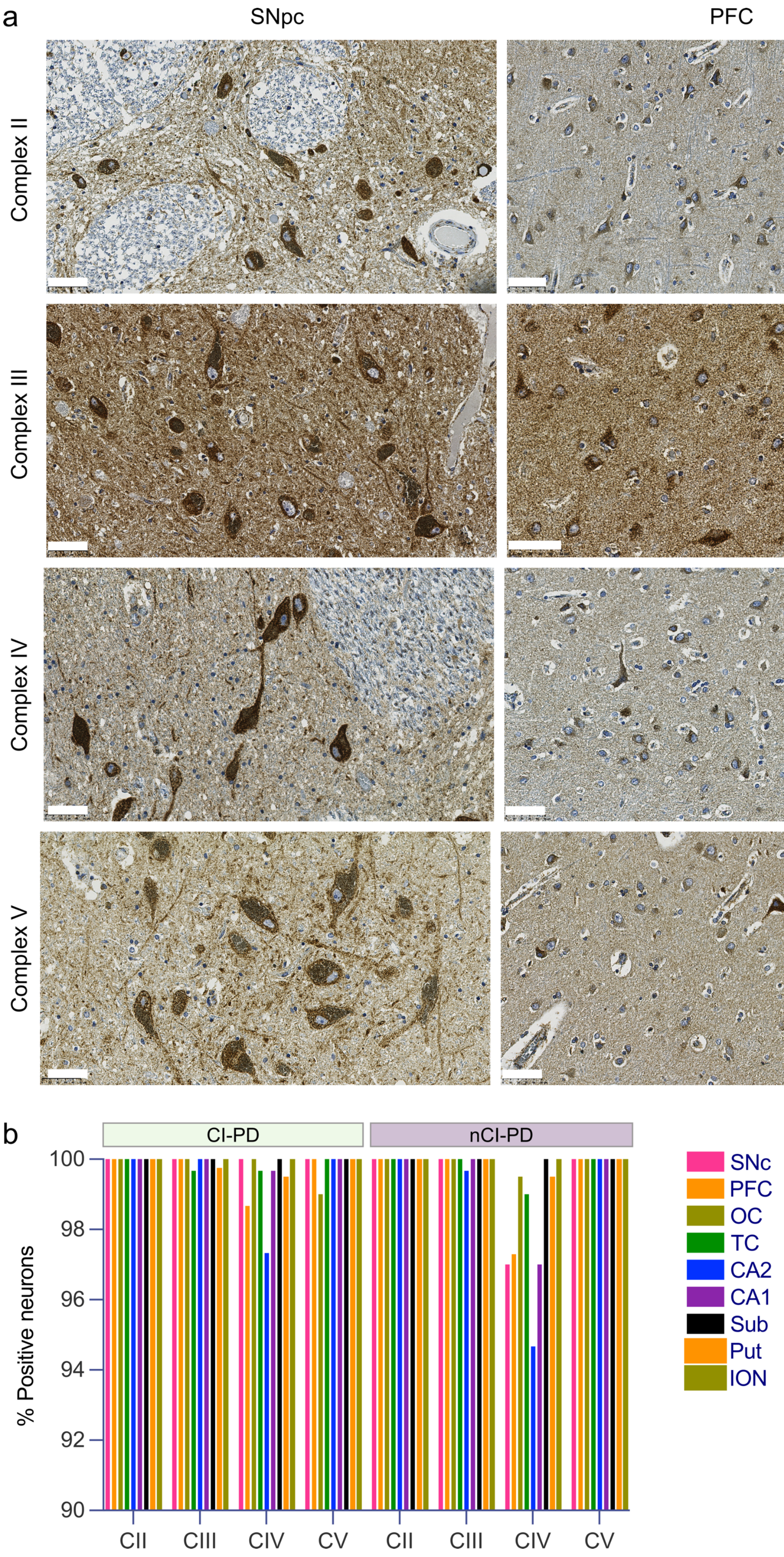
a
SNpc
PFC
Complex II
Complex III
Complex IV
Complex V
b
CI-PD
nCI-PD
% Positive neurons
100
98
96
94
92
90
CII
CIII
CIV
CV
CII
CIII
CIV
CV
SNc
PFC
OC
TC
CA2
CA1
Sub
Put
ION

**Fig. S2. Immunostaining for MRC complexes II-V**

(a) Representative images depicting immunostaining against complex II (SDHA), III (UQCRC2), IV (MTCOI), and V (ATP5A) in the SNpc and the PFC. Scalebar: 50 µm. (b) Plot showing the mean proportion (in %) of positive neurons from each of the nCI-PD and CI-PD-groups for each of the MRC complexes II (CI-PD: $n$ = 4, nCI-PD: $n$ = 3), III (CI-PD: $n$ = 4, nCI-PD: $n$ = 3), IV (CI-PD: $n$ = 3, nCI-PD: $n$ = 3) and V (CI-PD: $n$ = 3, nCI-PD: $n$ = 4). The number of included subjects from each region varied between two to four. The exact number of included subjects and the percentage positive neurons are given in Supplementary Data 3 and in the Source Data file. Neurons with complex II, III and V deficiencies are rare, while mild complex IV deficiency (generally ≤5% of the total regional cell count) was observed in both groups. CII: complex II, CIII: complex III, CIV: complex IV, CV: complex V, SNpc: substantia nigra pars compacta, PFC: prefrontal cortex, OC: occipital cortex, TC: temporal cortex, CA2: hippocampal region CA2, CA1: hippocampal region CA1, Sub: subiculum, Put: putamen, ION: inferior olivary nucleus

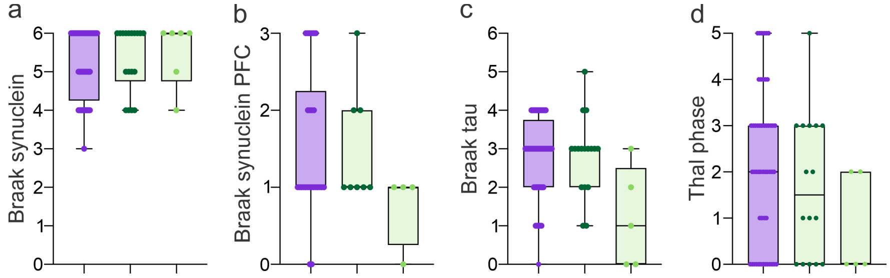


**Fig. S3. Neuropathology markers in CI-PD and nCIPD**

Box plots show individual values (dots), median and interquartile range (box) and minimum-maximum range (fences) of clinical data (y-axis) for the three iPD groups (nCI-PD: purple, CI-PD mild: dark green, CI-PD severe: light green). The number of included individuals is not equal in the different comparisons and is shown in Supplementary Data 5 and in the Source Data file. Statistical significance was tested between nCI-PD and the entire CI-PD group (i.e., not differentiating between mild and severe), due to the low number of individuals in the severe group, using the Mann-Whitney U-test. Pathology data were available from both cohorts. The exact P-values are given in Supplementary Data 5 and the Source Data file.

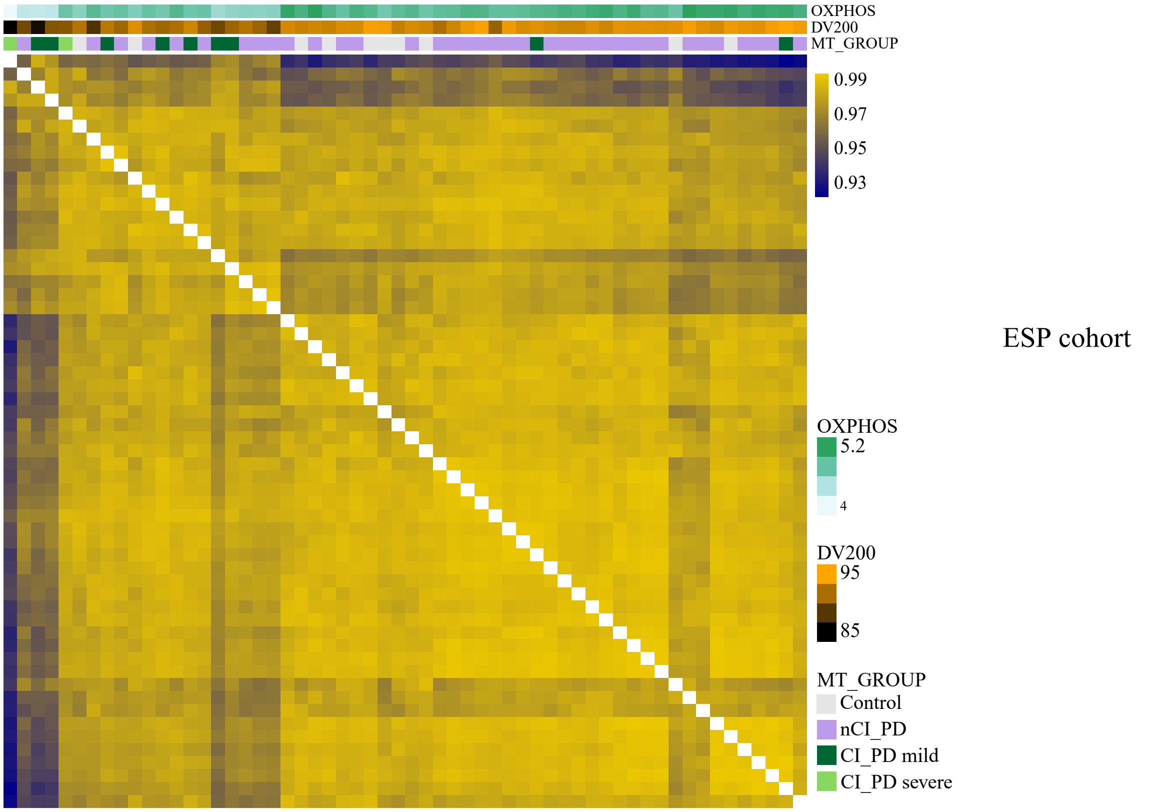


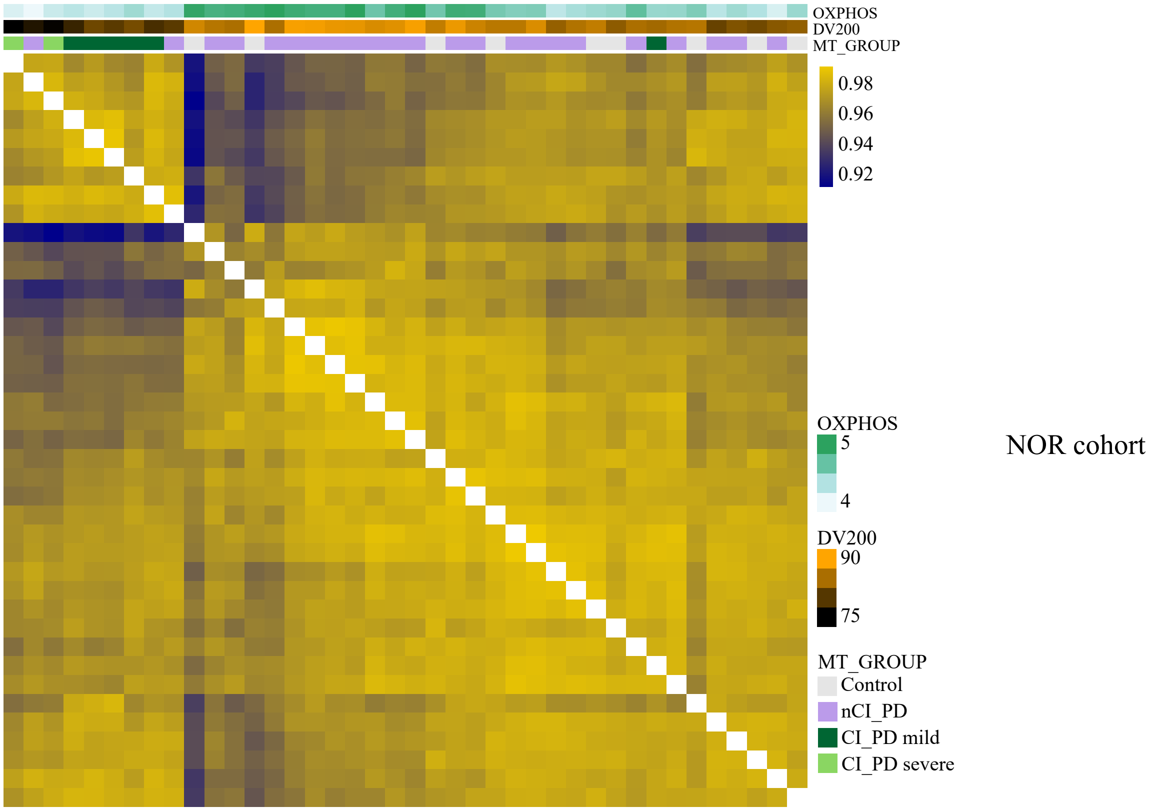


**Fig. S4. Sample-sample correlation of the transcriptomics data in NOR and ESP cohorts**

Hierarchical clustering of the samples based on sample-sample correlation in the ESP (top; $n = 58$) and NOR (bottom; $n = 40$) cohorts. The heatmap colors indicate Pearson's correlation values between each pair of samples. Subject demographics and experimental allocation are shown in Supplementary Data 1 and 2, respectively. OXPHOS: the first principal component of genes annotated to KEGG oxidative phosphorylation gene-set. MT_Group: group

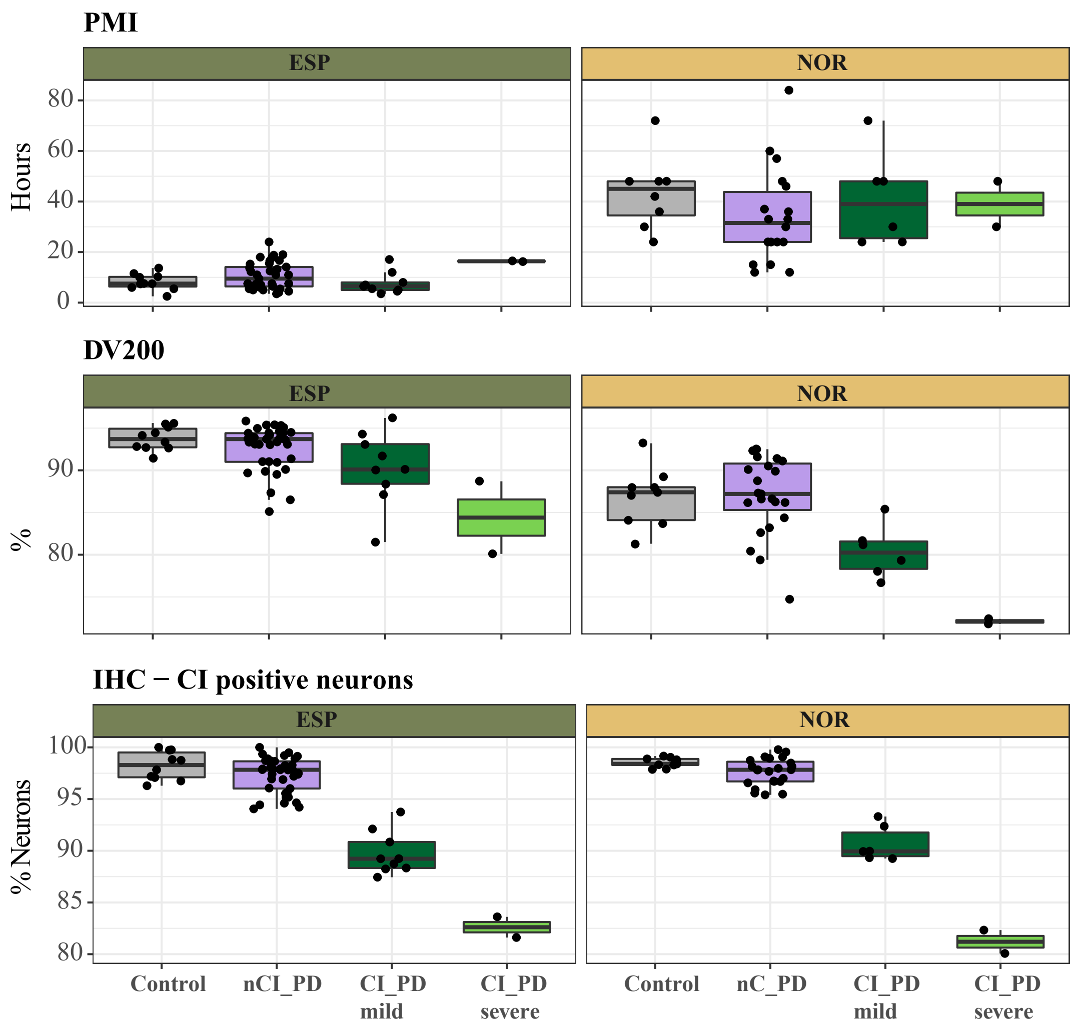


**Fig. S5. Lower DV200 is associated with the CI-PD subtype, but not with PMI**

In each of the NOR ($n$ = 40) and ESP ($n$ = 58) cohorts, the CI_PD subtypes were characterized by lower DV200 (y-axis, middle panel), compared both to controls and the nCI_ PD subtype. This could not be explained by PMI (y-axis, upper panel), since, in both cohorts, the PMI range of the CI_PD groups was similar to that of the controls (upper panel). The proportion of CI positive neurons (y-axis, lower panel) in the PFC of each group of subjects included in the RNA-seq analyses is shown in the bottom panel for comparison. Each point represents one individual. The boxplots represent the interquartile range (IQR) with the median indicated by the bold line. Whiskers are extensions from the top and the bottom of the boxplot by a value of 1.5*IQR. Subject demographics and experimental allocation are shown in Supplementary Data 1 and 2, respectively.

~Group + Sex + Age + PMI + Cohort (Model 2)

| | CI_PD | | nCI_PD | |
|---|---|---|---|---|
| | Group Coeficient (95%C.I.) | pValue | Group Coeficient (95%C.I.) | pValue |
| GabaVIPReln | −0.31 (−0.43, −0.18) | 4.7e−06 | −0.02 (−0.12, 0.08) | 0.65 |
| GabaRelnCalb | −0.29 (−0.41, −0.17) | 1e−05 | −0.02 (−0.11, 0.08) | 0.75 |
| Pyramidal | −0.26 (−0.38, −0.14) | 2.9e−05 | 0.03 (−0.06, 0.13) | 0.47 |
| Endothelial | 0.3 (0.16, 0.44) | 3.6e−05 | 0.06 (−0.05, 0.17) | 0.29 |
| OligoPrecursors | 0.28 (0.15, 0.4) | 3.9e−05 | 0 (−0.1, 0.11) | 0.93 |
| Astrocyte | 0.21 (0.11, 0.31) | 5.4e−05 | −0.02 (−0.1, 0.06) | 0.58 |
| GabaPV | −0.21 (−0.34, −0.09) | 0.0012 | 0.09 (−0.01, 0.19) | 0.095 |
| Microglia_deactivation | 0.16 (0.02, 0.29) | 0.023 | 0.01 (−0.1, 0.11) | 0.9 |
| Microglia | 0.15 (0.01, 0.29) | 0.031 | 0 (−0.11, 0.11) | 0.94 |
| Oligo | 0.1 (−0.01, 0.22) | 0.081 | 0.01 (−0.08, 0.1) | 0.8 |
| Microglia_activation | 0.11 (−0.03, 0.25) | 0.12 | −0.01 (−0.12, 0.1) | 0.82 |

Group Coefficient

0.2

0.0

−0.2

~Group + Sex + Age + PMI + Cohort + GabaVIPReln_MGP (Model 1)

| | CI_PD | | nCI_PD | |
|---|---|---|---|---|
| | Group Coeficient (95%C.I.) | pValue | Group Coeficient (95%C.I.) | pValue |
| Astrocyte | 0.09 (−0.01, 0.18) | 0.079 | −0.03 (−0.1, 0.04) | 0.36 |
| Endothelial | 0.11 (−0.02, 0.24) | 0.097 | 0.04 (−0.05, 0.14) | 0.33 |
| OligoPrecursors | 0.05 (−0.05, 0.15) | 0.29 | −0.01 (−0.08, 0.06) | 0.74 |
| GabaPV | 0.04 (−0.05, 0.12) | 0.37 | 0.1 (0.04, 0.16) | 0.00078 |
| Pyramidal | −0.04 (−0.12, 0.05) | 0.39 | 0.05 (−0.01, 0.11) | 0.095 |
| Oligo | −0.05 (−0.16, 0.06) | 0.39 | 0 (−0.08, 0.08) | 0.99 |
| GabaRelnCalb | −0.03 (−0.09, 0.04) | 0.45 | 0 (−0.04, 0.05) | 0.88 |
| Microglia_deactivation | 0.03 (−0.11, 0.16) | 0.72 | 0 (−0.1, 0.1) | 0.95 |
| Microglia_activation | −0.02 (−0.17, 0.12) | 0.75 | −0.02 (−0.13, 0.08) | 0.67 |
| Microglia | 0.02 (−0.12, 0.16) | 0.77 | −0.01 (−0.11, 0.1) | 0.91 |

**Fig. S6. Group differences in cell type MGPs**

The estimated group difference in MGPs of the cell types are shown. To determine the significance of the cell type MGPs, the cell types were added to a model in a step-wise regression manner. None of the cell type MGPs was significant in the nCI_PD group ($n$ = 60), based on the base model of the step-wise regression (Model_2). After adjustment for GabaVIPReln, none of the MGPs remained significant in the CI_PD group ($n$ = 19). The cell types are described in Mancarci et al. [60]. C.I.: confidence interval

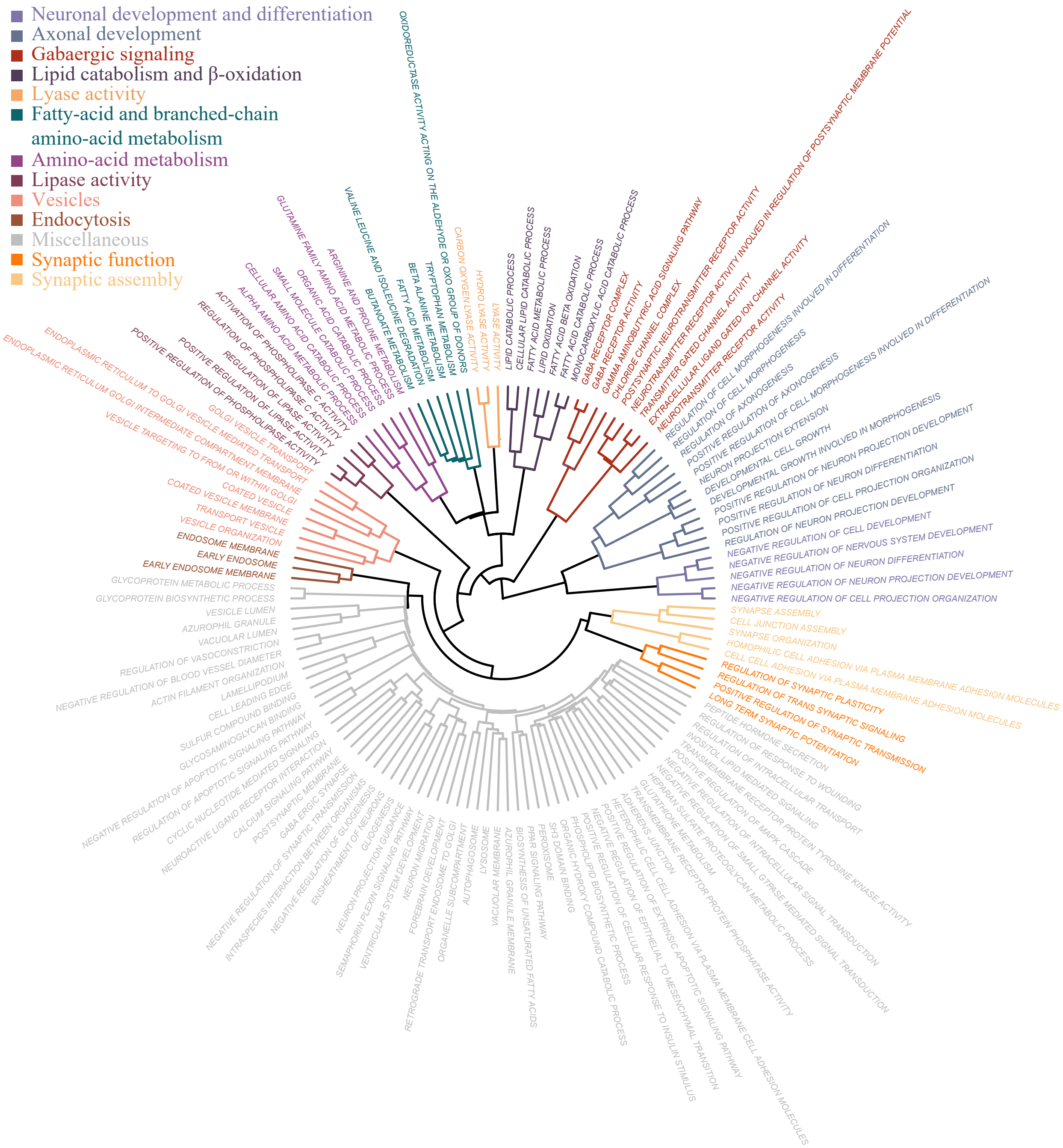


**Fig. S7. Significantly downregulated gene-sets in the nCI-PD group**

The enrichment analysis outcome of the nCI_PD group (*n* =60), based on the DE analysis using Model 2. Each leaf of the tree represents one significant gene-set. Jaccard similarity was used to cluster the gene-sets based on semantic similarity. Each cluster was given a name to represent the main function represented by its members.

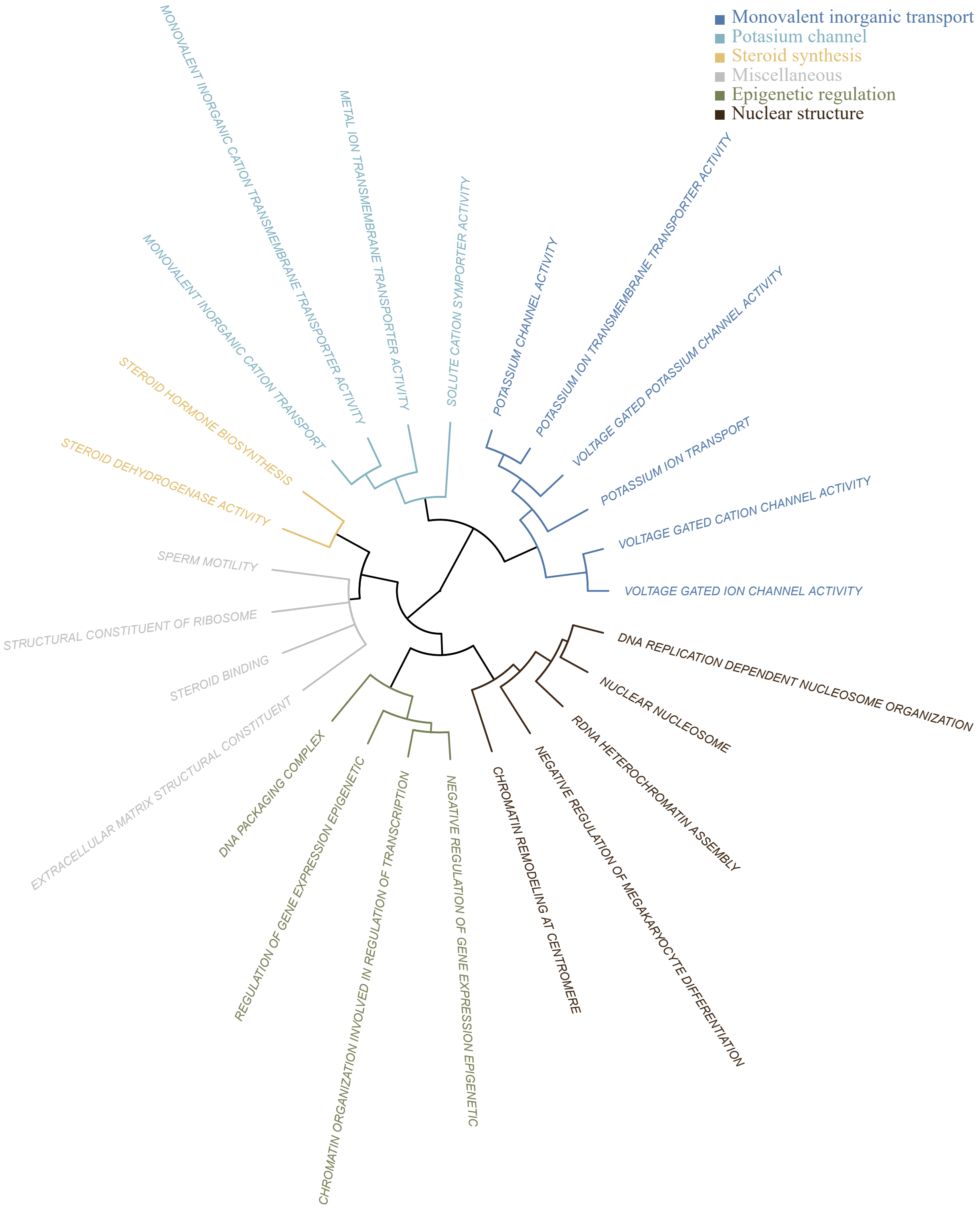


**Fig. S8. Significantly upregulated gene-sets in the nCI-PD group**

The enrichment analysis outcome of the nCI_PD group (*n* =60) based on DE analysis using Model_2. Each leaf of the tree represents one significant gene-set. Jaccard similarity was used to cluster the gene-sets based on semantic similarity. Each cluster was given a name to represent the main function represented by its members.

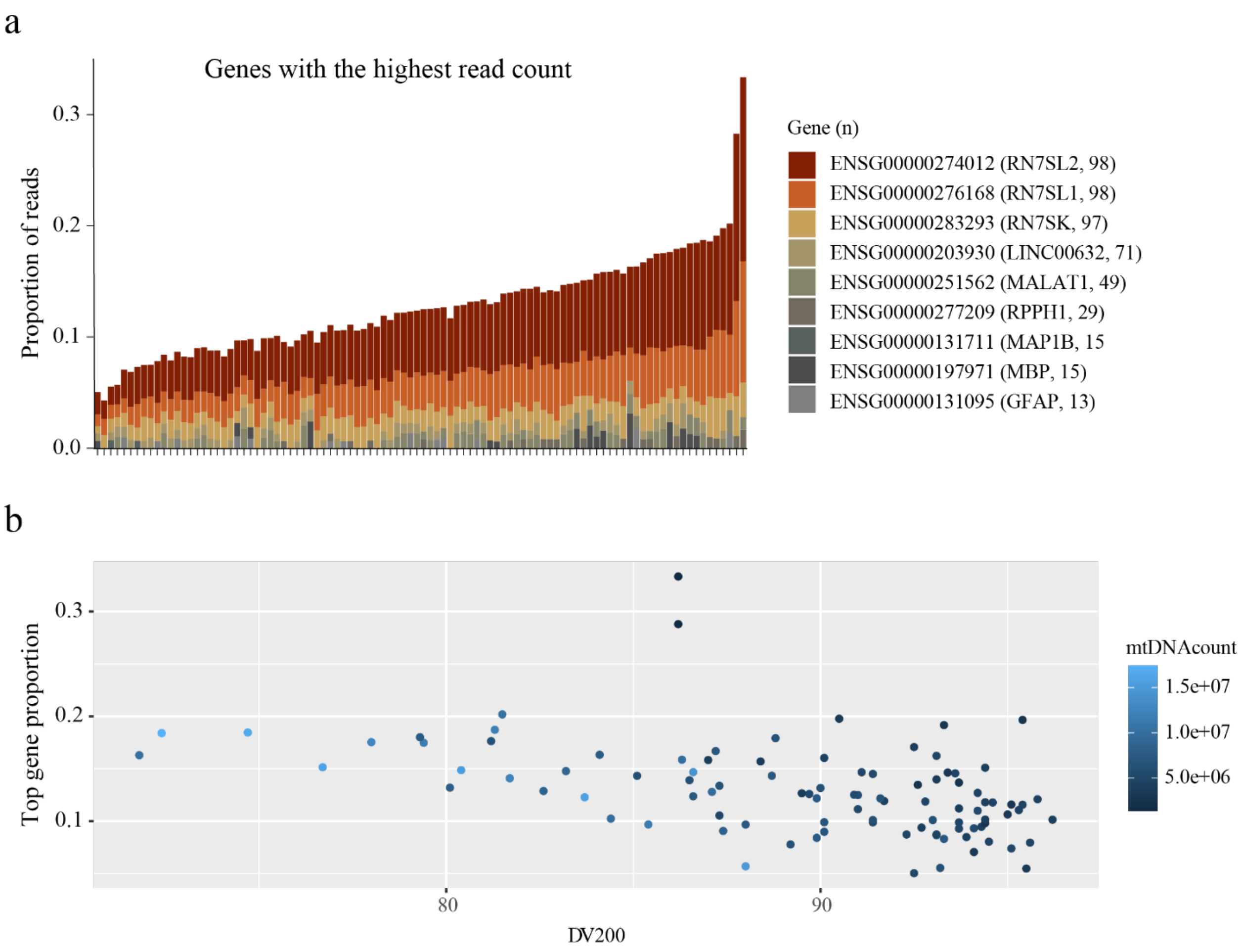


**Fig. S9 The highest expressed genes account for up to 34% of all reads in the RNA-seq data**
(a) The proportion of reads aligned to the five highest expressed genes in each sample (columns). n: the number of samples in which the gene was among the top five highest expressed. The three ribosomal genes – *RN7SL1*, *RN7SL2* and *RN7SK* were the highest expressed in all samples, accounting together to 5-32% of the total reads. Due to their impact on the total library size, and thus the downstream normalization steps, genes accounting for > 0.6% of the reads in more than 50% of the samples, were removed from downstream analyses. These included the three ribosomal genes, and *LINC00632.* (b) Correlation between DV200 (percent of fragments > 200 nucleotides) and the proportion of reads assigned to the top five highly expressed genes in a sample. Each point represents one sample. The small size of the three ribosomal genes, accounting for the vast majority of the top five highly expressed genes (299-331 bp), suggests that the variation in DV200

across the samples can be largely attributed to the variation in the proportion of these genes. mtDNAcount: the number of reads aligned to the mitochondrial genome.

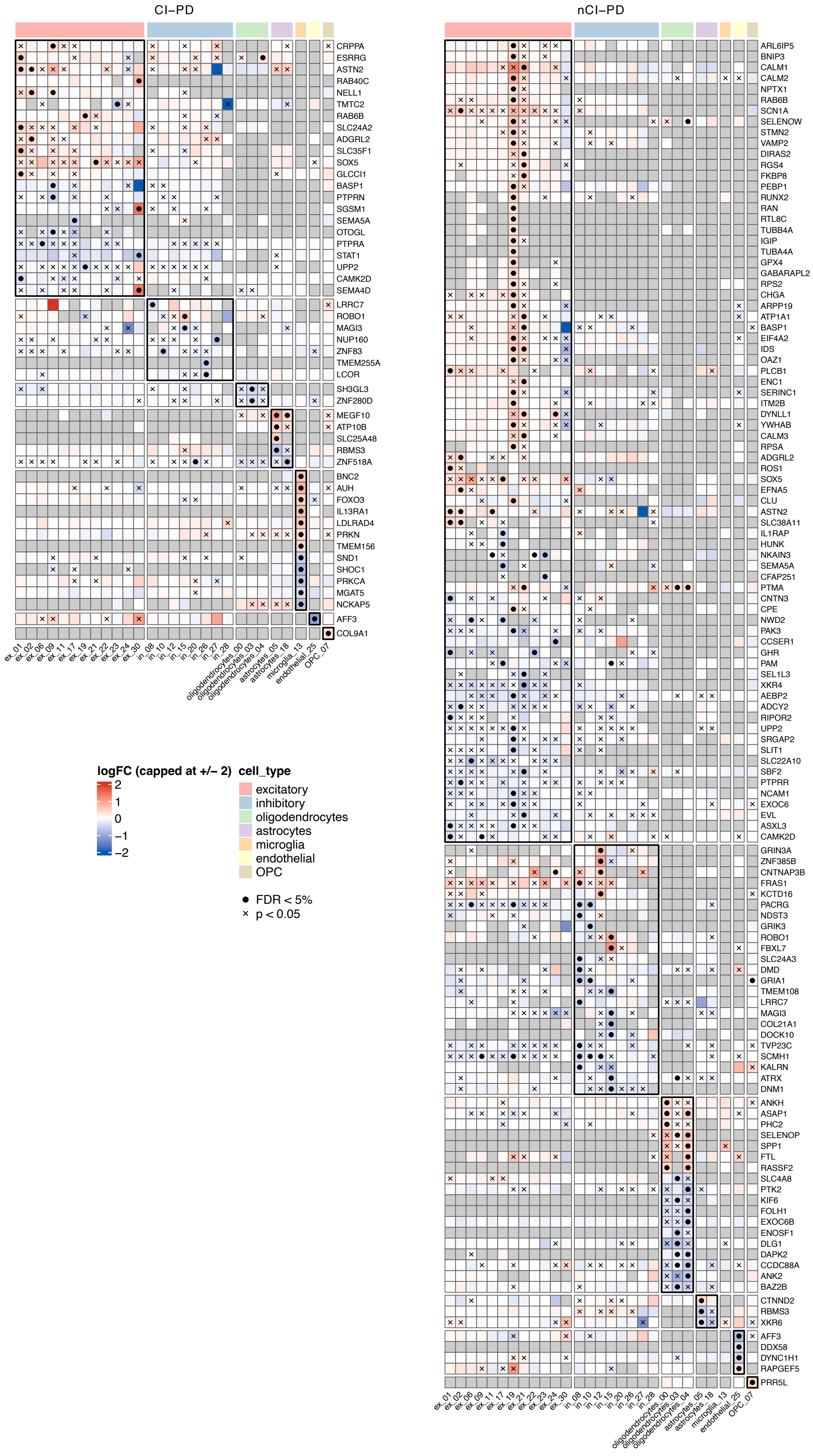
CI–PD
CRPPA
ESRRG
ASTN2
RAB40C
NELL1
TMTC2
RAB6B
SLC24A2
ADGRL2
SLC35F1
SOX5
GLCCI1
BASP1
PTPRN
SGSM1
SEMA5A
OTOGL
PTPRA
STAT1
UPP2
CAMK2D
SEMA4D
LRRC7
ROBO1
MAGI3
NUP160
ZNF83
TMEM255A
LCOR
SH3GL3
ZNF280D
MEGF10
ATP10B
SLC25A48
RBMS3
ZNF518A
BNC2
AUH
FOXO3
IL13RA1
LDLRAD4
PRKN
TMEM156
SND1
SHOC1
PRKCA
MGAT5
NCKAP5
AFF3
COL9A1
ex_01 ex_02 ex_06 ex_09 ex_11 ex_17 ex_19 ex_21 ex_22 ex_23 ex_24 ex_30 in_08 in_10 in_12 in_15 in_20 in_26 in_27 in_28 oligodendrocytes_00 oligodendrocytes_03 oligodendrocytes_04 astrocytes_05 astrocytes_18 microglia_13 endothelial_25 OPC_07
nCI–PD
ARL6IP5
BNIP3
CALM1
CALM2
NPTX1
RAB6B
SCN1A
SELENOW
STMN2
VAMP2
DIRAS2
RGS4
FKBP8
PEBP1
RUNX2
RAN
RTL8C
TUBB4A
IGIP
TUBA4A
GPX4
GABARAPL2
RPS2
CHGA
ARPP19
ATP1A1
BASP1
EIF4A2
IDS
OAZ1
PLCB1
ENC1
SERINC1
ITM2B
DYNLL1
YWHAB
CALM3
RPSA
ADGRL2
ROS1
SOX5
EFNA5
CLU
ASTN2
SLC38A11
IL1RAP
HUNK
NKAIN3
SEMA5A
CFAP251
PTMA
CNTN3
CPE
NWD2
PAK3
CCSER1
GHR
PAM
SEL1L3
XKR4
AEBP2
ADCY2
RIPOR2
UPP2
SRGAP2
SLIT1
SLC22A10
SBF2
PTPRR
NCAM1
EXOC6
EVL
ASXL3
CAMK2D
GRIN3A
ZNF385B
CNTNAP3B
FRAS1
KCTD16
PACRG
NDST3
GRIK3
ROBO1
FBXL7
SLC24A3
DMD
GRIA1
TMEM108
LRRC7
MAGI3
COL21A1
DOCK10
TVP23C
SCMH1
KALRN
ATRX
DNM1
ANKH
ASAP1
PHC2
SELENOP
SPP1
FTL
RASSF2
SLC4A8
PTK2
KIF6
FOLH1
EXOC6B
ENOSF1
DLG1
DAPK2
CCDC88A
ANK2
BAZ2B
CTNND2
RBMS3
XKR6
AFF3
DDX58
DYNC1H1
RAPGEF5
PRR5L
ex_01 ex_02 ex_06 ex_09 ex_11 ex_17 ex_19 ex_21 ex_22 ex_23 ex_24 ex_30 in_08 in_10 in_12 in_15 in_20 in_26 in_27 in_28 oligodendrocytes_00 oligodendrocytes_03 oligodendrocytes_04 astrocytes_05 astrocytes_18 microglia_13 endothelial_25 OPC_07
logFC (capped at +/– 2)
2
1
0
–1
–2
cell_type
excitatory
inhibitory
oligodendrocytes
astrocytes
microglia
endothelial
OPC
● FDR < 5%
× p < 0.05

111

**Fig. S10. Differentially expressed transcripts in snRNA-seq**

Genes that were differentially expressed (at FDR < 5% with minimum fold change of 25%) in at least one of the cell type clusters are represented in the rows of the heatmaps. Only autosomal protein-coding genes are represented. Columns correspond to cell type clusters, grouped by main cortical cell types (ex: excitatory neurons, in: inhibitory neurons, oligodendrocytes astrocytes, microglia, endothelial cells, and oligodendrocyte precursor cells; OPC). Cell colors correspond to the estimated log2-fold change in expression with respect to controls (red for upregulation and blue for downregulation). Grey cells correspond to genes that did not reach the required expression threshold in the cell type cluster. Crosses represent nominal significance in the cluster ($P < 0.05$) while black dots indicate statistical significance at FDR < 5%). Left-hand side heatmap displays the results for the CI-PD ($n = 7$) *vs* controls ($n = 6$) contrast, right-hand side for the nCI-PD ($n = 5$) vs controls.

125

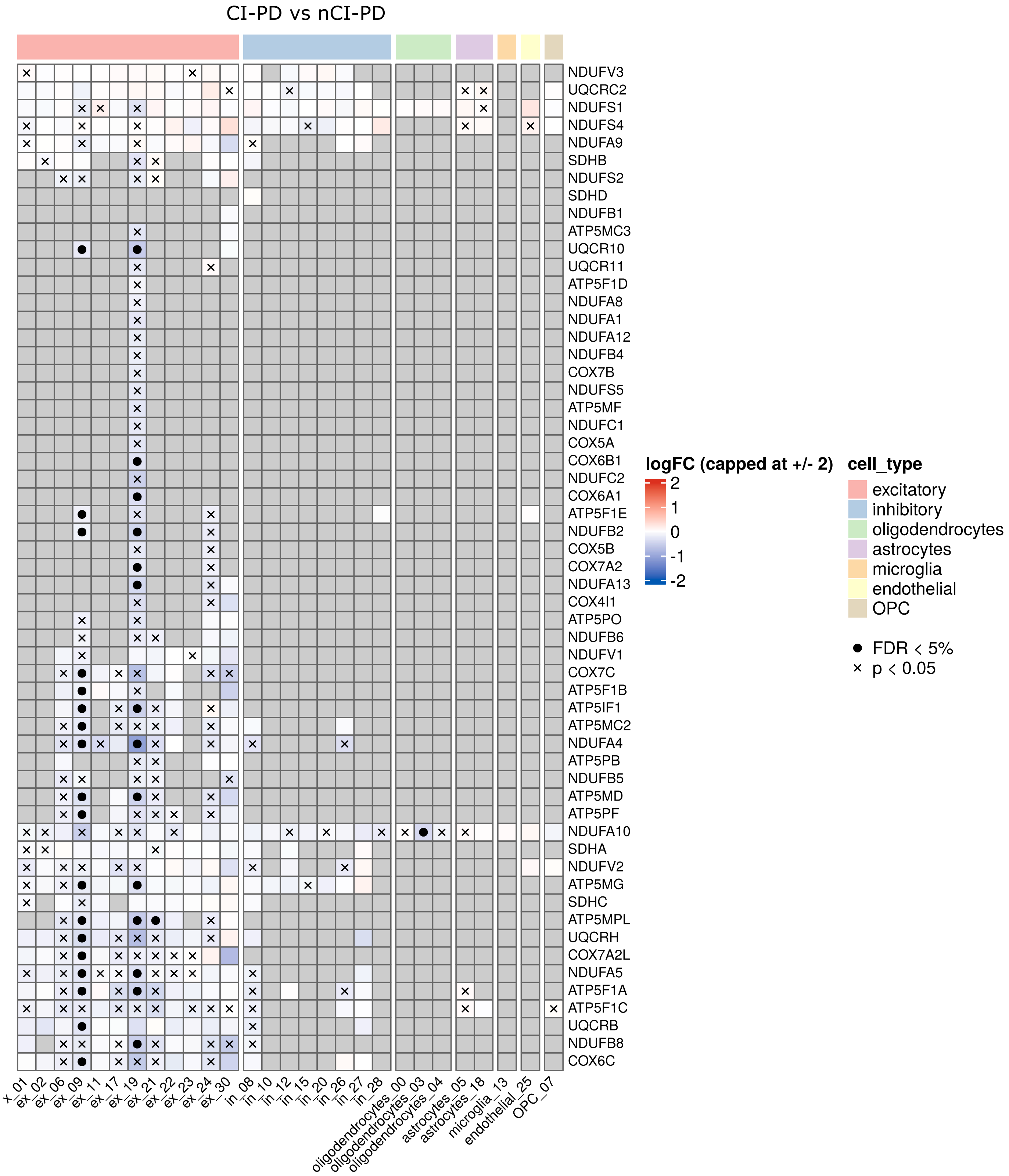


**Fig. S11. Differentially expressed MRC transcripts in snRNA-seq: CI-PD vs nCI-PD**

The heatmap shows results for the contrast of CI-PD (*n* = 7) vs nCI-PD (*n* = 5). Only autosomal protein-coding MRC genes are represented. Columns correspond to cell type clusters, grouped by main cortical cell types (ex: excitatory neurons, in: inhibitory neurons, oligodendrocytes astrocytes, microglia, endothelial cells, and oligodendrocyte precursor cells; OPC). Cell colors correspond to the estimated log2-fold change in expression with respect to controls (red for upregulation and blue for downregulation). Grey cells correspond to genes that did not reach the required expression threshold in the cell type cluster. Crosses represent nominal significance in the cluster ($P < 0.05$) while black dots indicate statistical significance (FDR < 5%).

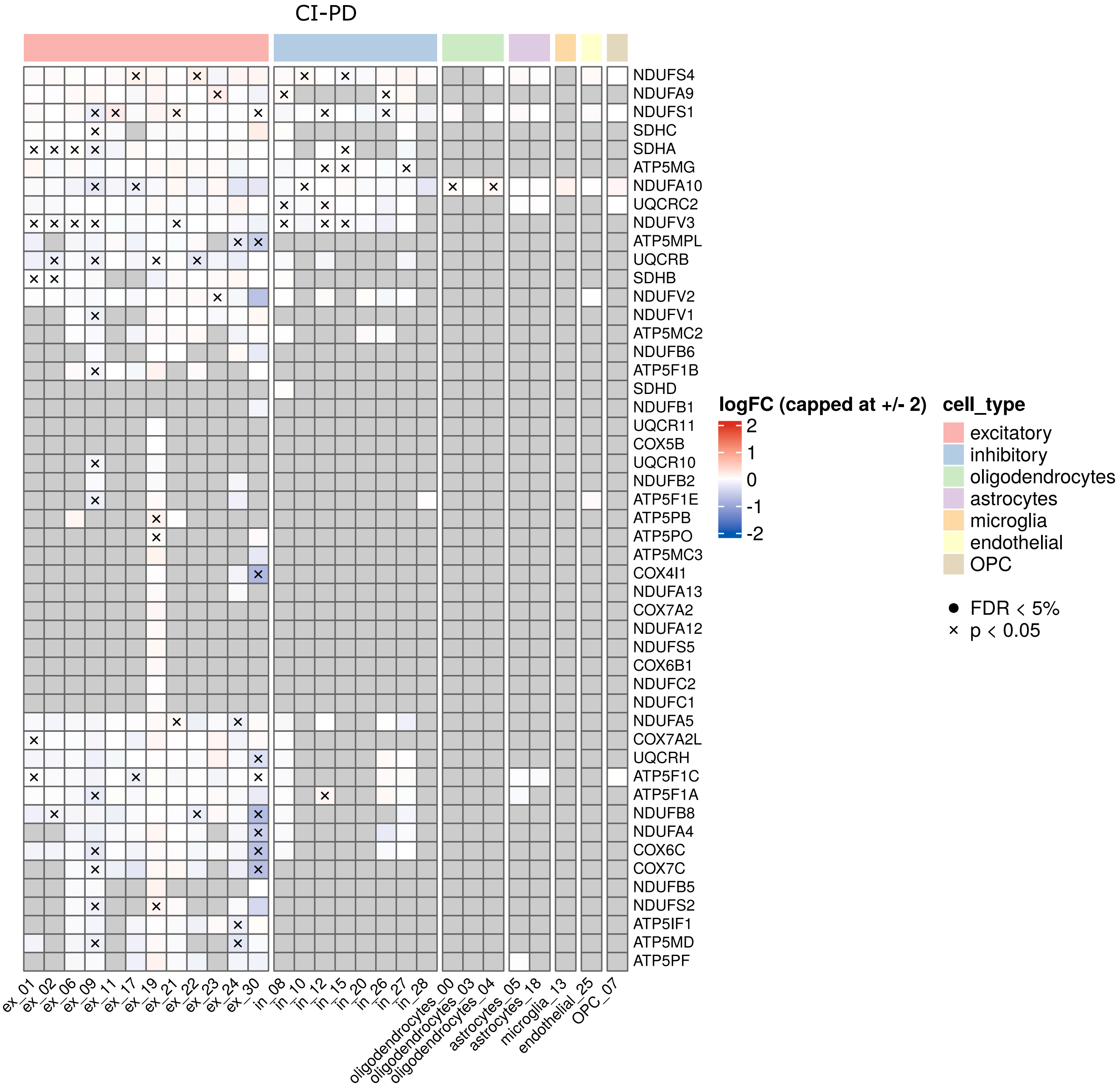


**Fig. S12. Differentially expressed MRC transcripts in snRNA-seq: CI-PD vs controls**

The heatmap shows results for the CI-PD group ($n$ = 7) vs controls ($n$ = 6) as a contrast. Only autosomal protein-coding MRC genes are represented. Columns correspond to cell type clusters, grouped by main cortical cell types (ex: excitatory neurons, in: inhibitory neurons, oligodendrocytes astrocytes, microglia, endothelial cells, and oligodendrocyte precursor cells; OPC). Cell colors correspond to the estimated log2-fold change in expression with respect to controls (red for upregulation and blue for downregulation). Grey cells correspond to genes that did not reach the required expression threshold in the cell type cluster. Crosses represent nominal significance in the cluster ($P < 0.05$) while black dots indicate statistical significance ($FDR < 5\%$).

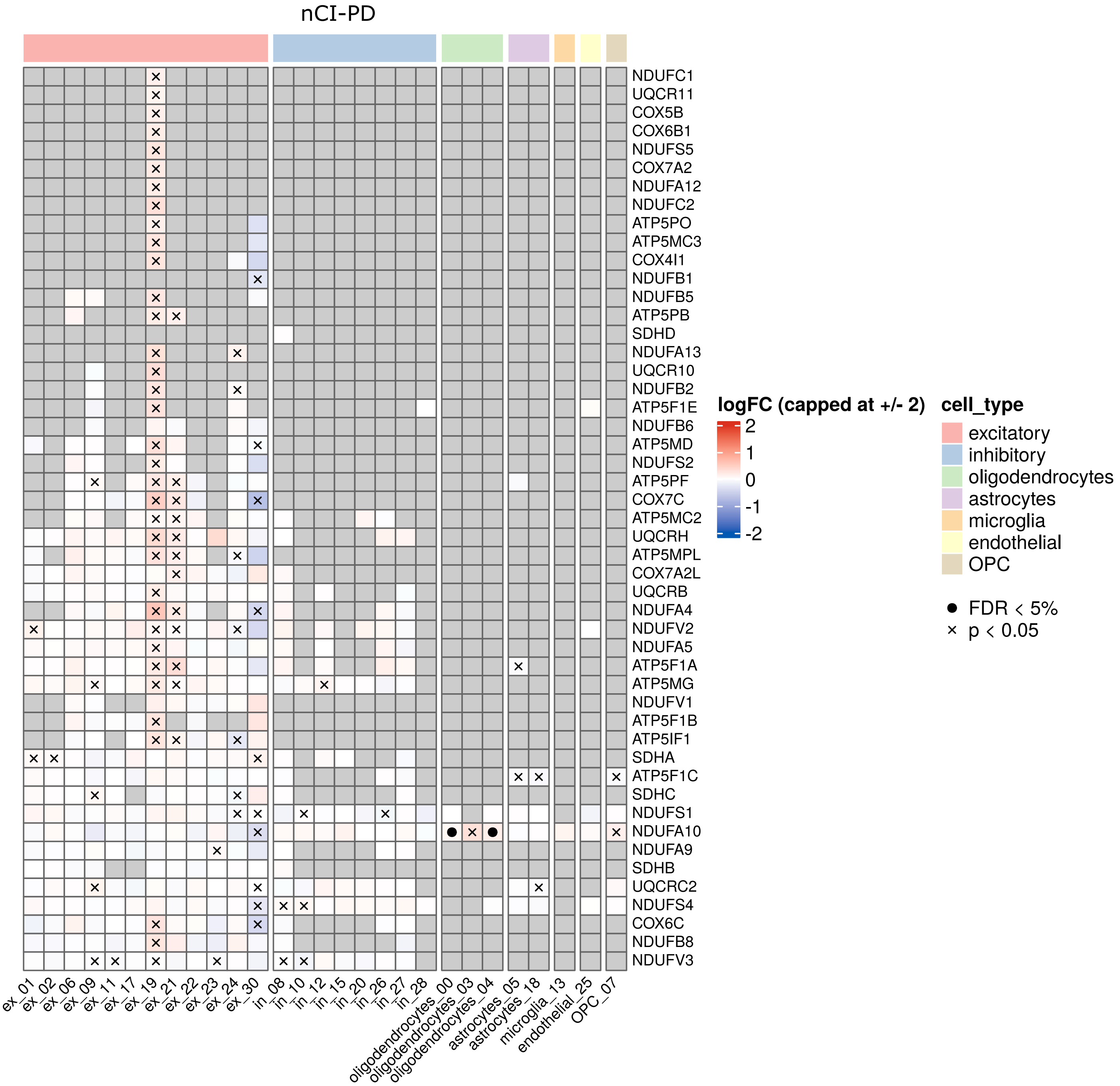


**Fig. S13. Differentially expressed MRC transcripts in snRNA-seq: nCI-PD vs controls**

The heatmap shows results for the nCI-PD group ($n$ = 5) vs controls ($n$ = 6) as a contrast. Only autosomal protein-coding MRC genes are represented. Columns correspond to cell type clusters, grouped by main cortical cell types (ex: excitatory neurons, in: inhibitory neurons, oligodendrocytes astrocytes, microglia, endothelial cells, and oligodendrocyte precursor cells;

OPC). Cell colors correspond to the estimated log2-fold change in expression with respect to controls (red for upregulation and blue for downregulation). Grey cells correspond to genes that did not reach the required expression threshold in the cell type cluster. Crosses represent nominal significance in the cluster ($P < 0.05$) while black dots indicate statistical significance (FDR < 5%).

**Supplementary Data 1. (Separate file; Excel).**

**Supplementary Data 2. (Separate file; Excel).**

**Supplementary Data 3. (Separate file; Excel).**

**Supplementary Data 4. (Separate file; Excel).**

**Supplementary Data 5. (Separate file; Excel).**

**Supplementary Data 6. (Separate file; Excel).**

**Supplementary Data 7. (Separate file; Excel).**

**Supplementary Data 8. (Separate file; Excel).**

**Supplementary Data 9. (Separate file; Excel).**

**Supplementary Data 10. (Separate file; Excel).**

**Supplementary Data 11. (Separate file; Excel)**

**Supplementary Data 12. (Separate file; Excel)**

**Supplementary Data 13. (Separate file; Excel)**
