## Supplementary material for "Mitochondrial complex I deficiency stratifies idiopathic Parkinson’s disease": Inventory of supplementary information

**Inventory of Supporting Information**

**Supplementary Figure S1.**

Immunostaining of CI subunits in the SNpc

**Supplementary Figure S2.**

Immunostaining for MRC complexes II-V

**Supplementary Figure S3.**

Neuropathology markers in CI-PD and nCIPD

**Supplementary Figure S4.**

Sample-sample correlation of the transcriptomics data in NOR and ESP cohorts

**Supplementary Figure S5.**

Lower DV200 is associated with the CI-PD subtype, but not with PMI

**Supplementary Figure S6.**

Group differences in cell type MGPs

**Supplementary Figure S7.**

Significantly downregulated gene-sets in the nCI-PD group

**Supplementary Figure S8.**

Significantly upregulated gene-sets in the nCI-PD group

**Supplementary Figure S9.**

The highest expressed genes account for up to 34% of all reads in the RNA-seq data

**Supplementary Figure S10.**

Differentially expressed transcripts in snRNA-seq

**Supplementary Figure S11.**

Differentially expressed MRC transcripts in snRNA-seq: CI-PD vs nCI-PD

**Supplementary Figure S12.**

Differentially expressed MRC transcripts in snRNA-seq: CI-PD vs controls

**Supplementary Figure S13.**

Differentially expressed MRC transcripts in snRNA-seq: nCI-PD vs controls

**Supplementary Data 1.**

Subject demographics, pathology scores and clinical data.

**Supplementary Data 2.**

Sample experiment allocation.

**Supplementary Data 3.**

MRC IHC data.

**Supplementary Data 4.**

Data and statistics for IHC analysis of CI (NDUFS4) in 17 brain regions.

**Supplementary Data 5.**

Data and statistics for clinical and pathological parameters.

**Supplementary Data 6.**

Data and statistics for single neuron mtDNA analyses.

**Supplementary Data 7.**

Transcriptomics. Differential gene expression analysis comparing each of the nCI-PD and CI-PD groups to controls in the combined cohort. Results of all three models (Model 1-3) are shown.

**Supplementary Data 8.**

Transcriptomics. Pathway-enrichment analysis for each of the nCI-PD and CI-PD groups in the combined cohort. Results of all three models (Model 1-3) are shown.

**Supplementary Data 9.**

Single-nuclei RNA-seq. Differential gene expression analysis for each cell type, between CI-PD and controls.

**Supplementary Data 10.**

Single-nuclei RNA-seq. Differential gene expression analysis for each cell type, between nCI-PD and controls.

**Supplementary Data 11.**

Single-nuclei RNA-seq. Differential gene expression analysis for each cell type, between CI-PD and nCI-PD.

**Supplementary Data 12.**

Single-nuclei RNA-seq. Functional enrichment of differentially expressed genes for each cell type cluster by overrepresentation analysis of the Kyoto Encyclopedia of Genes and Genomes (KEGG) pathway.

**Supplementary Data 13.**

Single-nuclei RNA-seq. Overrepresentation analysis on the union of significantly differentially expressed genes in excitatory neurons, inhibitory neurons and glia
